# Identification of a MarR subfamily that regulates arsenic resistance genes

**DOI:** 10.1101/2021.08.12.456183

**Authors:** Yanshuang Yu, Renwei Feng, Jichen Chen, Yuanping Li, Jinxuan Liang, Zhenchen Xie, Hend A. Alwathnani, Barry P. Rosen, Anne Grove, Jian Chen, Christopher Rensing

## Abstract

Members of the family of Multiple Antibiotic Resistance Regulators (MarR) often regulate genes that encode antibiotic resistance in bacteria. In this study, comprehensive analyses were performed to determine the function of an atypical MarR homolog in *Achromobacter* sp. As-55. Genomic analyses showed that this *marR* is located in an arsenic gene island in *Achromobacter* sp. As-55 adjacent to an *arsV* gene. ArsV is a flavin-dependent monooxygenase that confers resistance to the antibiotic methylarsenite (MAs(III)), the organoarsenic compound roxarsone(III) (Rox(III)), and the inorganic antimonite (Sb(III)). Similar *marR* genes are widely distributed in arsenic-resistant bacteria. Phylogenetic analyses showed that these MarRs are found in operons predicted to be involved in resistance to inorganic and organic arsenic species, so the subfamily was named MarR_ars_. MarR_ars_ orthologs have three conserved cysteine residues, which are Cys36, Cys37 and Cys157 in *Achromobacter* sp. As-55, mutation of which compromises the response to MAs(III)/Sb(III). GFP-fluorescent biosensor assays show that AdMarR_ars_ (MarR protein of *Achromobacter deleyi* As-55) responds to trivalent As(III) and Sb(III) but not to pentavalent As(V) or Sb(V). The results of RT-qPCR assays show that *arsV* is expressed constitutively in a *marR* deletion mutant, indicating that *marR* represses transcription of *arsV*. Moreover, electrophoretic mobility shift assays (EMSA) demonstrate that AdMarR_ars_ binds to the promoters of both *marR* and *arsV* in the absence of ligands and that DNA binding is relieved upon binding of As(III) and Sb(III). Our results demonstrate that AdMarR_ars_ is a novel As(III)/Sb(III)-responsive transcriptional repressor that controls expression of *arsV*, which confers resistance to MAs(III), Roxarsone(III) and Sb(III). AdMarR_ars_ and its orthologs form a subfamily of MarR proteins that regulate genes conferring resistance to arsenic-containing antibiotics.

**IMPORTANCE:** In this study, a MarR family member, AdMarR_ars_ was shown to regulate the *arsV* gene, which confers resistance to arsenic-containing antibiotics. It is a founding member of a distinct subfamily that we refer to as MarR_ars_, regulating genes conferring resistance to arsenic and antimony antibiotic compounds. AdMarR_ars_ was shown to be a repressor containing conserved cysteine residues that are required to bind As(III) and Sb(III), leading to a conformational change and subsequent derepression. Here we show that members of the MarR family are involved in regulating arsenic-containing compounds.

---

Arsenic and antimony pollution have attracted considerable attention in recent years due to their adverse effect on the environment and human health (1). Although As and Sb pollution poses a health threat to humans, animals and plants, some microorganisms survive in environments with high concentrations of these metalloids, and even utilize them for growth. These microbes have adapted metabolic pathways that incorporate As and Sb or evolved mechanisms to confer resistance or detoxify them, thus playing a substantial role in the metalloid biogeochemical cycle (2, 3).

Active efflux of As and Sb out of the cytoplasm is the most common mechanism of metalloid resistance in bacteria (4). Such resistance is encoded in various *ars* operons found in many species of bacteria. These *ars* operons are carried on plasmids and chromosomes, and their expression is usually induced by As(III) and Sb(III) (5). Among proteins encoded on *ars* operons, ArsR was the first identified member of the family of ArsR/SmtB transcriptional repressors, regulating its expression and downstream *ars* genes. In the absence of arsenic, homodimeric ArsR binds to the promoter region of the operon to repress *ars* gene expression. In the presence of inducers such as As(III), Sb(III) (6-8) or MAs(III) (9), ArsR undergoes a conformational change and dissociates from the promoter DNA sequence, leading to expression of the operon. The most common *ars* operons contain an *arsR* gene, an *arsC* gene encoding an As(V) reductase and a gene encoding an As(III) efflux permease, usually either *arsB* or *acr3* (6-8, 10). Additional common genes include *arsA* encoding an arsenic ATPase subunit ArsA (11) and *arsD* encoding an arsenite chaperone that delivers As(III) to the ArsAB transporter complex (12). Additional less common *ars* genes have been discovered, including *arsH* encoding an organoarsenical oxidase (13), *arsI* encoding an MAs(III) demethylase (14), *arsN*, which encodes an N-acetyltranferase that confers resistance to the arsenic antibiotic arsinothricin (15), *arsO* encoding a putative flavin-binding monooxygenase (16), *arsJ*, which confers arsenate resistance together with GADPH (glyceraldehyde-3-phosphate dehydrogenase) (17), *arsP* encoding a methylarsenite efflux permease (18), *arsV* encoding a NADPH-dependent flavin monooxygenase (19), MacAB encoding an ABC-type efflux protein (20) and ars*TX* conferring functions related to thioredoxin metabolism (21). It is likely that more *ars* genes await discovery. MarR, first identified in *Escherichia coli*, is a transcriptional repressor unrelated to ArsR (22, 23). It is a multi-antibiotic-resistance regulator that functions as a homodimer, with a characteristic winged helix-turn-helix (wHTH) as the DNA binding motif in addition to a ligand-binding region (24). In the absence of inducer, MarR binds to its own gene promoter, repressing transcription of itself as well downstream genes organized in the same operon. After binding the ligand, MarR dissociates from the promoter region, enabling DNA transcription (25). MarR family proteins regulate functions involving antibiotic resistance and handling of oxidative stress, virulence factor production, catabolism of aromatic compounds and as a master regulator in bacteria such as *Burkholderia* sp. (25). In addition, MarRs regulate catabolism of lignin and other substances and synthesis of antibiotics (26).

Some MarR family members interact with metals. The transcription of the zinc transporter operon (*zit*) in *Lactococcus lactis* is regulated by the MarR family transcriptional regulator ZitR. When the zinc concentration in the environment is too high, Zn(II) binds to ZitR and changes the conformation of ZitR to tightly bind to the *zit* promoter, thereby inhibiting the transcription of the *zit* operon and subsequently reducing transport of Zn into cells. Under Zn starvation, unliganded ZitR dissociates from the *zit* promoter sequence and relieves inhibition of the *zit* operon (27, 28). *E. coli* MarR is a prototypical member of the MarR family. MarR either directly regulates one or two genes encoding a specific function or regulates a single operon such as *marRAB*. MarA then functions as an activator of many genes involved in a pleiotropic response. Cu(II) oxidizes a unique cysteine residue (Cys80 in *E. coli* MarR) in its DNA-binding domain, forming a disulfide bond between two MarR dimers, producing a conformational change that renders it unable to bind to the *marRAB* promoter, thereby derepressing expression of the *marRAB* operon (29).

To date, As/Sb has not been shown to regulate any member of the MarR family. In this study, we analyzed the genome of the highly arsenite-resistant bacterium *Achromobacter deleyi* As-55 (MIC 36 mM) (GenBank accession number: CP074375.1), which was isolated from antimony mine in Lengshuijiang, Hunan Province, China. We identified and characterized a *marR* gene adjacent to an *ars*/*aio* operon and analyzed the As-55 AdMarR_ars_ protein at the molecular and genetic levels.

## RESULTS AND DISCUSSION

### Genes for MarR orthologs are widely distributed in *ars* operons

Examination of the *Achromobacter* sp. As-55 genome identified a *marR*-like gene near an *aio/ars* operon. Genes annotated as a *marR* were also located in predicted *ars* operons of other arsenic resistant bacteria (Fig. 1). These *marR* genes are present upstream of an *arsV* gene in *Paenibacillus* sp. NC1, *Roseateles aquatilis, Ktedonobacter racemifer* DSM 44963, *Luteitalea pratensis* and *Eoetvoesia caeni*; or located upstream of an *arsN* gene in *Duganella* sp. CF458, *Chloroflexi bacterium* 54-19, *Ktedonobacter racemifer* DSM 44963, *Deinococcus* sp. YIM 77859 and *Tahibacter aquaticus*. This association suggests that this particular type of MarR regulates expression of *arsV, arsP* and *arsN*. These MarR proteins form a distinct subfamily within the MarR family that we renamed for clarity as MarR_ars_ (Fig. 2). Interestingly, MarR_ars_ is predicted to regulate expression of genes that have previously been shown to confer resistance to the arsenic-containing antibiotics such as MAs(III) and arsenothricin (15, 30), in keeping with the overall role of MarRs as regulators of antibiotic resistance (24).

**FIG 1.**
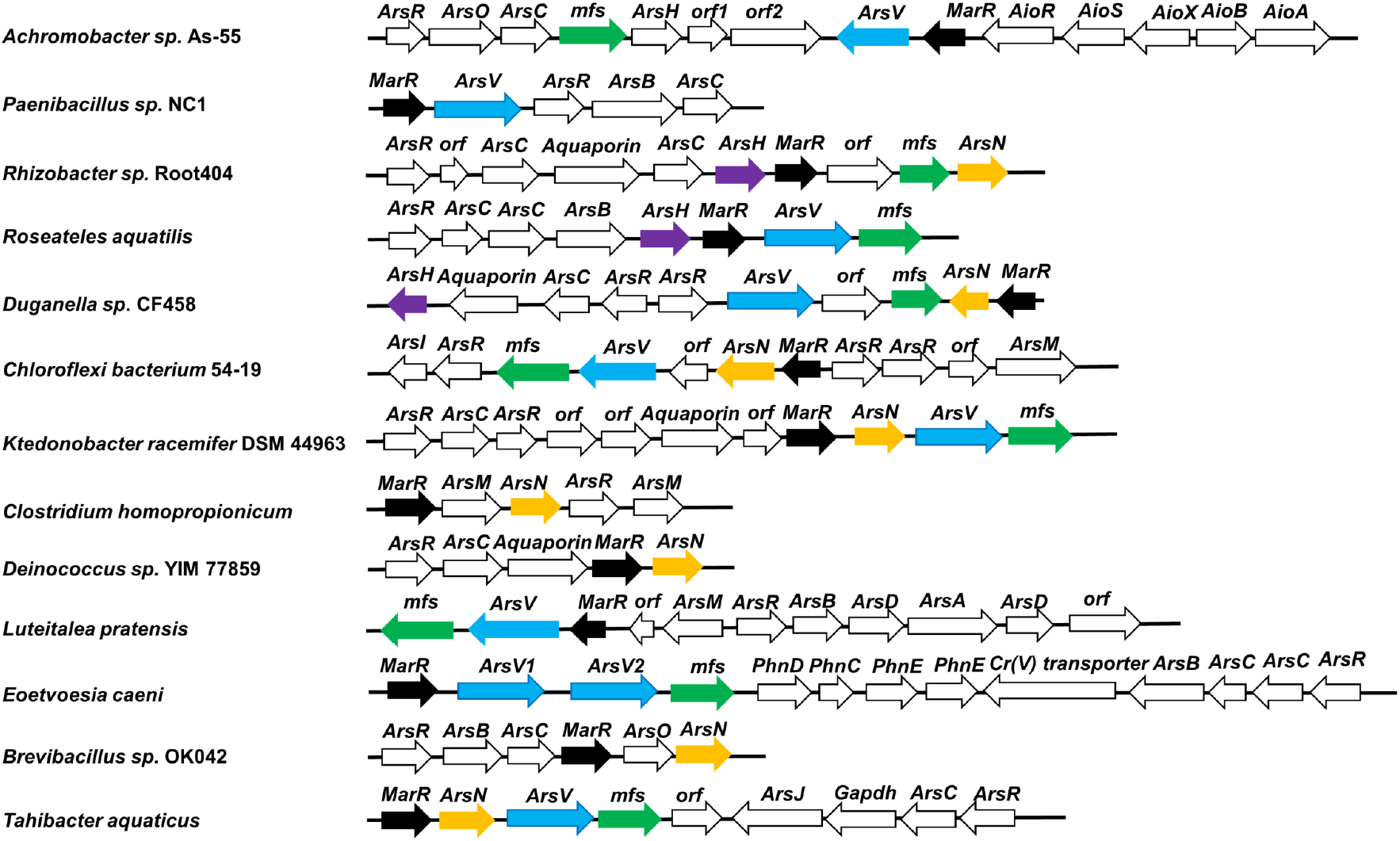
*marR*_*ars*_ genes distributed in *ars* operons from different organisms. Shown are representative *ars* operons (accession numbers in parentheses) containing *marR* genes (black fill). *Achromobacter* sp. As-55 (NZ_CP074375.1), *Paenibacillus* sp. NC1 (NZ_QEVW01000012.1), *Rhizobacter* sp. Root404 (NZ_LMDS01000005.1), *Roseateles aquatilis* (NZ_NIOF01000003.1), *Duganella* sp. CF458 (NZ_FOOF01000005.1), *Chloroflexi bacterium* 54-19 (MKTJ01000032.1), *Ktedonobacter racemifer* DSM 44963 (NZ_ADVG01000002.1), *Clostridium homopropionicum* (NZ_FOOL01000002.1), *Deinococcus* sp. YIM 77859 (NZ_JQNI01000004.1), *Luteitalea pratensis* (NZ_CP015136.1), *Eoetvoesia caeni* (NZ_JACCEU010000005.1), *Brevibacillus* sp. OK042 (NZ_FORT01000019.1), *Tahibacter aquaticus* (NZ_SNZH01000010.1).

**FIG 2.**
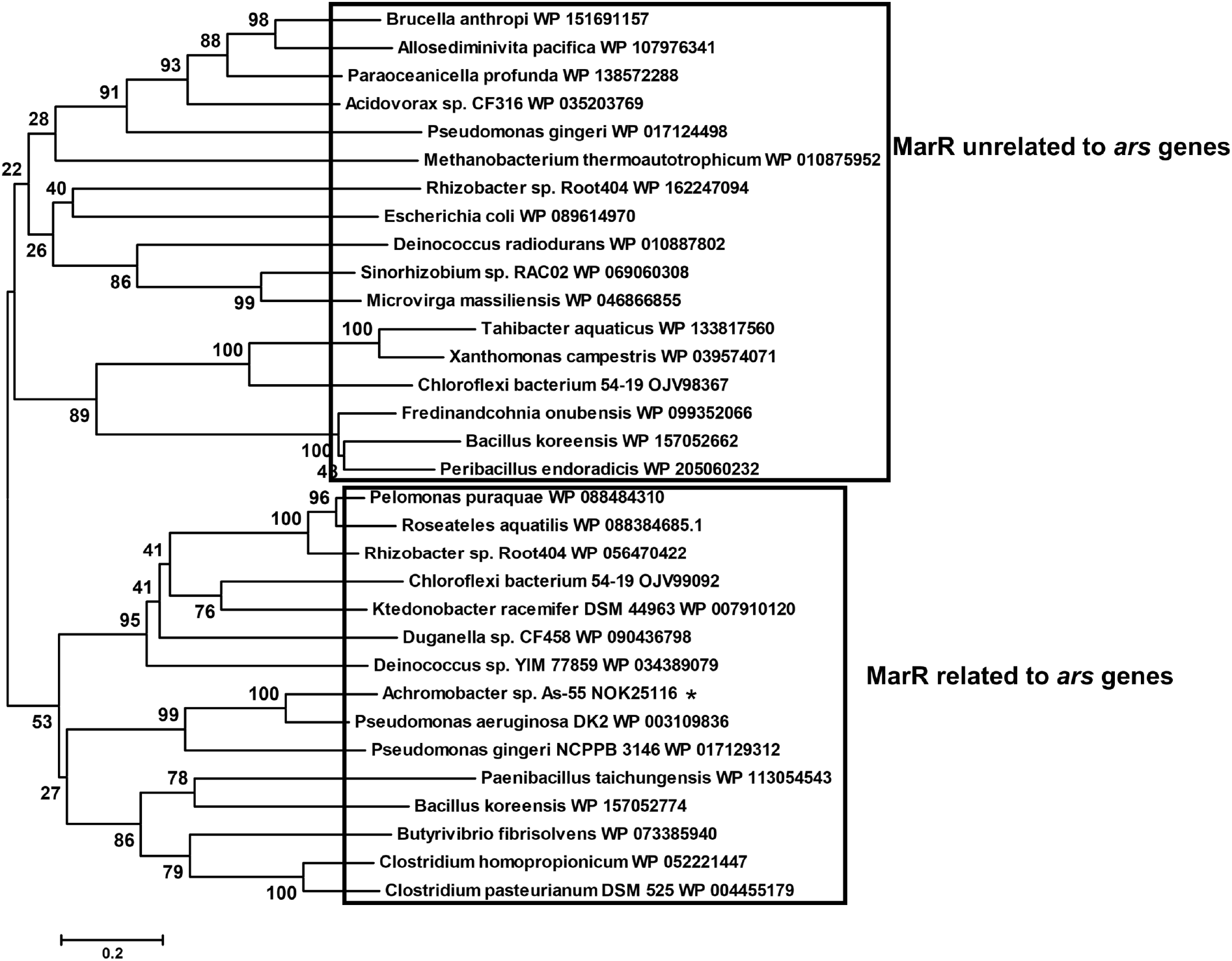
The neighbor-joining phylogenetic tree constructed based on MarR proteins from different bacteria. AdMarR_ars_ from *Achromobacter* sp. As-55 is indicated with an asterisk. It was constructed using the neighbor-joining algorithm with MEGA 6.0. The bootstrap values (based on 1, 000 replications) are indicated at the branch nodes. GenBank protein accession numbers are shown in parentheses after each protein. The bar represents 0.2 amino acid substitution per site.

### AdMarR_ars_ is an As(III)/Sb(III)-responsive transcriptional repressor

A *marR*_*ars*_ deletion mutant (Δ*marR*_*ars*_) in *Achromobacter* sp. As-55 was generated, and the expression of the adjacent *arsV* gene was examined by RT-qPCR (Fig. 3). Expression of *arsV* was up-regulated in wild-type cells by 0.2 and 2 mM As(III) or 0.05 and 0.2 mM Sb(III) compared to the wild type As-55 with no addition of metalloids. In Δ*marR*_*ars*_ cells, *arsV* was highly expressed even in the absence of metalloids, demonstrating that AdMarR_*ars*_ functions as a repressor of *arsV* and that the metalloids serve as inducers of AdMarR_ars_. Consistent with constitutively high expression of *arsV* in Δ*marR*_*ars*_ cells, the *marR*_*ars*_ deletion conferred resistance to roxarsone (Fig. 4). Rox(III) was shown to be much more toxic than Rox(V). Wild type cells were unable to grow in 16 μM Rox(III), while cells of *A. deleyi* Δ*marR*_*ars*_ grew in 16 μM Rox(III). While both strains were able to grow at 1.6 mM Rox(V) (not shown), only the *A. deleyi* Δ*marR*_*ars*_ grew in 3.2 mM Rox(V). It is conceivable that at high concentrations of Rox(V), small amounts of Rox(V) were reduced to Rox(III), thereby generating toxicity. As noted above, *arsV* is predicted to encode a flavin-dependent oxidoreductase oxidizing MAs(III) (19). Other genes in the vicinity of marR_ars_ were not to be regulated by MarR_ars_ indicating only *marR* and *arsV* were regulated by MarR_ars_ (Fig. S1 and S2)

**FIG 3.**
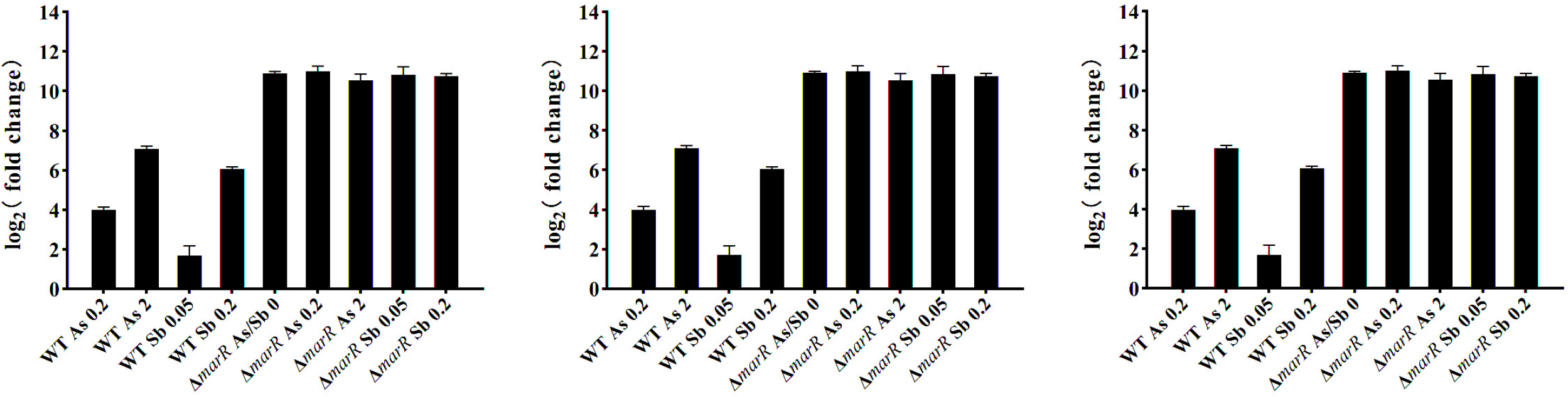
Expression of *arsV* of *Achromobacter* sp. As-55 (WT) and *marR*_*ars*_ mutant (Δ*marR*_*ars*_) under As(III)/Sb(III) exposure. WT As 0.2/As 2: *Achromobacter* sp. As-55 under 0.2/2 mM As(III) exposure; WT Sb 0.05/Sb 0.2: *Achromobacter* sp. As-55 under 0.05/0.2 mM Sb(III) exposure; Δ*marR*_*ars*_ As/Sb 0: *marR*_*ars*_ mutant without metal added; Δ*marR*_*ars*_ As 0.2/As 2: *marR*_*ars*_ mutant under 0.2/2 mM As(III) exposure; Δ*marR*_*ars*_ As 0.2/As 2: *marR*_*ars*_ mutant under 0.05/0.2 mM Sb(III) exposure. The log_2_(fold change) is reported relative to treatment of *Achromobacter* sp. As-55 (WT) with no metals added.

**FIG 4.**
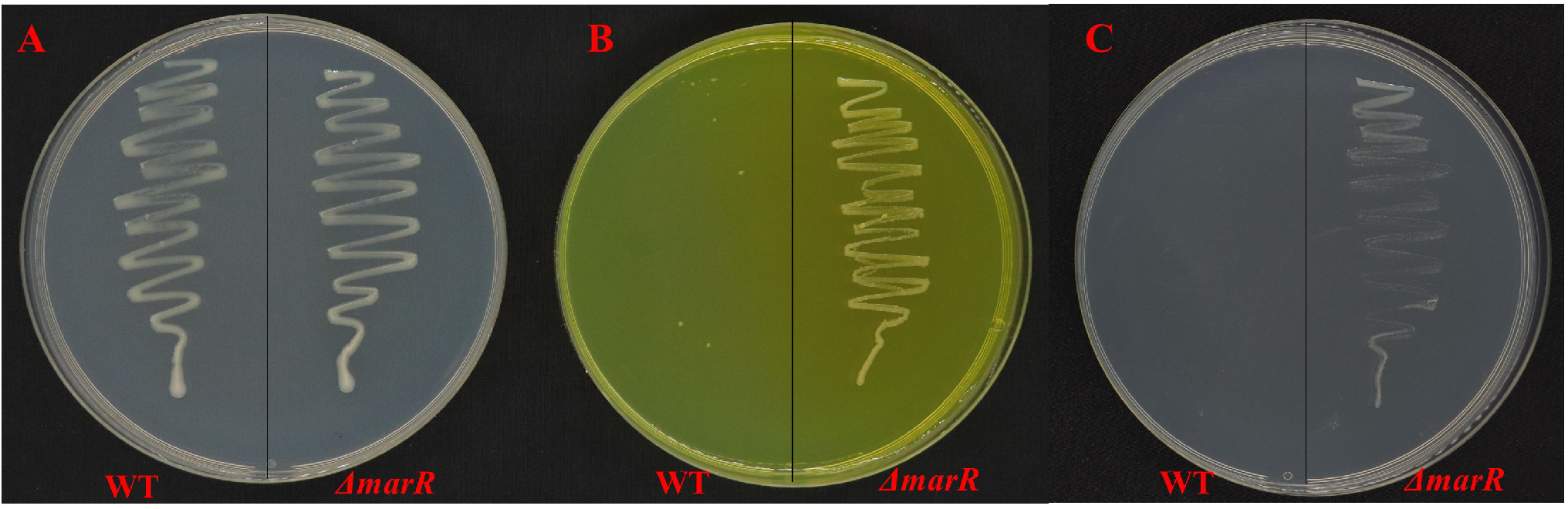
Deletion of AdmarR_ars_ confers resistance to roxarsone. (A) Growth of WT and *marR*_*ars*_ mutant (Δ*marR*_*ars*_) in absence of roxarsone. Δ*marR*_*ars*_ displays resistance to Rox(V) (B) and Rox(III) (C) compared to wild type *Achromobacter* sp. As-55 (WT). The overnight culture was streaked on R2A solid medium containing 3.2 mM Rox(V) (B), 16 µM Rox(III) (C) and no roxarsone as control.

### ArsV confers resistance to organoarsenicals

To examine the function of the *Achromobacter sp*. As-55 *arsV* gene product, the gene was cloned into plasmid pTOPO, constructing plasmid pTOPO-*arsV* with *arsV* expressed under the universal Km promoter, which was expressed in the arsenic-sensitive *E. coli* strain AW3110*Δars* (31). Metalloid resistance was assayed by measuring growth and reporting OD_600_ after one day exposure to the indicated compounds. The strain containing pTOPO-*arsV* grew well in lysogeny broth (LB) medium containing 16 μM MAs(III), 4-8 mM Rox(V) or 8 μM Rox(III), while the strain containing the vector did not grow under the same conditions (Fig. 5), demonstrating that ArsV confers resistance to MAs(III), Rox(III) and Rox(V).

**FIG 5.**
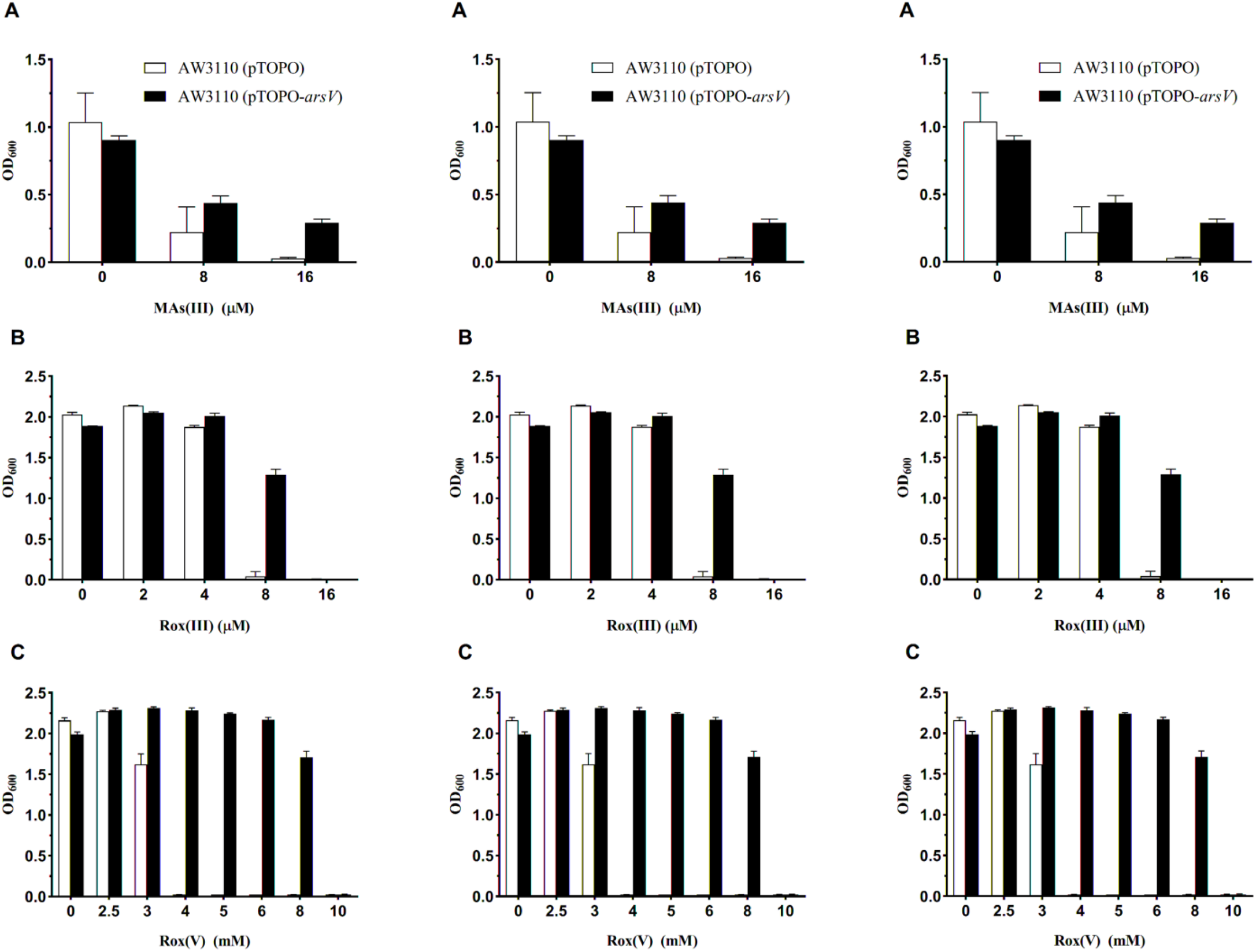
ArsV conferred resistance to MAs(III), Rox(III) and Rox(V) in *E. coli* AW3110. Growth of *E. coli* AW3110 containing plasmid pTOPO or pTOPO-*arsV* was measured after the addition of different concentrations of MAs(III) (A), Rox(III) (B), Rox(V) (C) in liquid LB medium. The data is the average of three independent replicates with standard deviation.

### AdMarR_ars_ is derepressed by metalloids

We hypothesized that AdMarR_*ars*_ is autoregulatory and controls expression of *arsV*. Electrophoretic mobility shift assays (EMSA) were used to examine the interaction between AdMarR_ars_ and the regulatory DNA encompassing the non-coding region but also a small part of the coding region up and downstream of the *arsV* and *marR*_*ars*_ promoters. Purified AdMarR_ars_ was incubated with either Cy5.5-labeled *marR* promoter or *arsV* promoter, and electrophoretic mobility of the DNA-protein complexes were retarded compared to the free probe (Fig. 6). With increasing As(III) and Sb(III) concentrations, the electrophoretic shifts of the Cy5.5-labeled probes were gradually reduced, suggesting AdMarR_ars_ regulating expression of its own gene and *arsV* in a metalloid-dependent manner.

**FIG 6.**
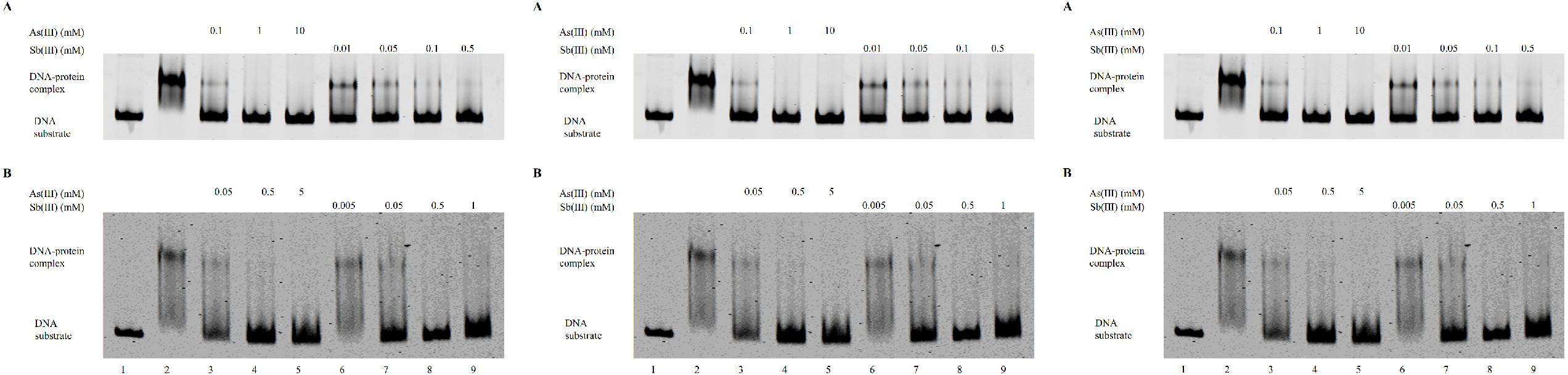
AdMarR_ars_ binds *arsV* and *marR*_*ars*_ promoters. EMSA assays with *arsV* promoter (A) and *marR*_*ars*_ promoter (B). Lanes 1: Cy5.5-labeled *arsV*/*marR*_*ars*_ promoter probe without AdMarR_ars_ protein; Lanes 2: Cy5.5-labeled *arsV*/*marR*_*ars*_ promoter probe with MarR protein; Lanes 3-5: Cy5.5-labeled *arsV*/*marR*_*ars*_ promoter probe with AdMarR_ars_protein co-incubated with various amount of As(III); Lanes 6-9: Cy5.5-labeled *arsV*/*marR*_*ars*_ promoter probe with AdMarR_ars_ protein co-incubated with various amount of Sb(III). Representative of three replicates.

### Role of conserved cysteine residues in the MarR_ars_ subfamily

A neighbor-joining phylogenetic tree was constructed based on MarR_ars_ proteins from different bacteria (Fig. 2). Here, we selected genes encoding MarR that were part of an *ars* operon and selected other representative members of the MarR family of regulators not involved in arsenic resistance. These putative MarR_ars_ repressors form a distinct subfamily within the MarR family. A multiple sequence alignment of these MarR_ars_ regulatory proteins shows that three cysteine residues (Cys36, Cys37 and Cys157) are conserved in AdMarR_ars_ (Fig. S3). We predict that these are involved in As(III)/Sb(III) binding, since cysteine triads generally bind As(III) and Sb(III) in ArsR repressors (32), although their location in the primary sequences of the proteins vary (Fig. S4). A homology model of AdMarR_ars_ constructed using the MarR structure from *Methanosarcina mazei* Go1 (PDB ID: 3S2W) (https://www.rcsb.org/structure/3S2W) as a template indicates that they could form an As(III)/Sb(III) binding site in the folded repressor (Fig. S5). To examine the role of the conserved cysteine residues in MarR_ars_ function, Cys36, Cys37 and Cys157 were individually altered to serine residues by site-directed mutagenesis. We used a GFP biosensor strain (33) in which *AdmarR*_*ars*_ is under control of the *ara* promoter and *gfp* is under control of the *AdmarR*_*ars*_ promoter; in cells expressing the C36S, C37S, C157S AdMarR_ars_ variants, *gfp* expression was compared with cells expressing wild type AdMarR_ars_ following exposure to 0, 10, 20, 30 or 40 µM As(III) (Fig. 7A). The fluorescence intensity increased with increasing concentrations of As(III). The fluorescence intensity of the cells expressing wild type AdMarR_ars_ was much higher than the three mutants, consistent with a loss of As(III) binding by the mutants. In addition, AdMarR_ars_ responded to As(III) and Sb(III), but not to As(V) or Sb(V) (Fig. 7B). These findings indicate thiolate-dependent binding due to the soft-metal character of both As(III) and Sb(III) and in analogy to ArsR specificity would be predicted to be achieved by resulting conformational change not by affinity.

**FIG 7.**
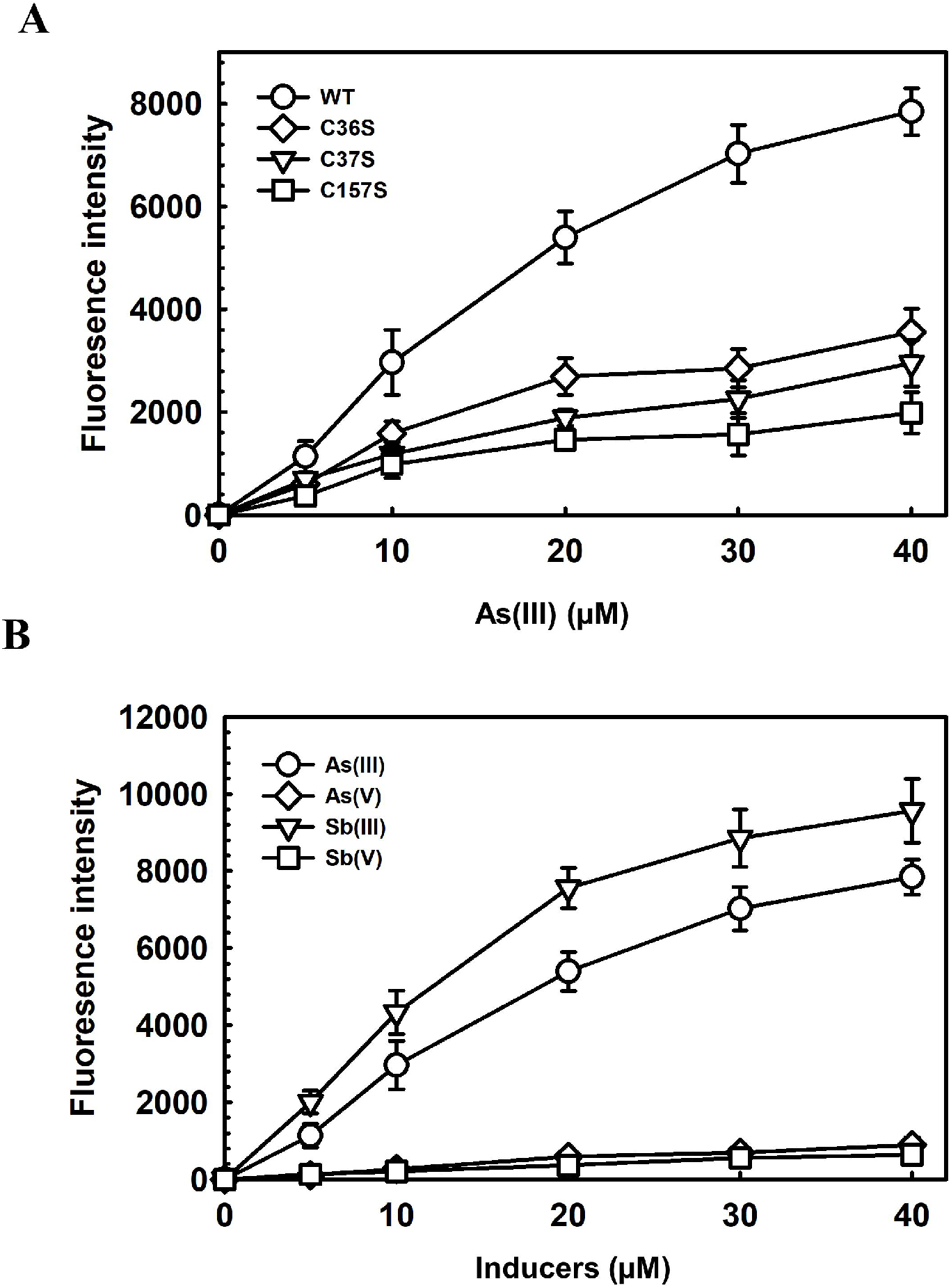
(A) Binding of As(III) to AsMarR_ars_ involves specific cysteine residues. Expression of the *gfp* reporter gene was assayed as described in materials and methods. GFP induction of cysteine mutants (C36S, C37S, C157S) and AdMarR_ars_ (WT) with increasing concentration of As(III). (B) GFP induction with different inducers of AdMarR_ars_ in *Achromobacter* sp. As-55. Comparison of the response of the bacterial biosensor to arsenic and antimony.

### Conclusions

The results of this study support our hypothesis that AdMarR_ars_ is an As(III)/Sb(III)-responsive transcriptional regulator. It regulates genes that confer resistance to the antibiotic MAs(III) such as *arsV*, which encodes a flavin-dependent monooxygenase that oxidizes highly toxic MAs(III) to relatively nontoxic MAs(V). Genes for MarR_ars_ orthologs are widely distributed in bacteria, indicating that the regulatory function mediated by MarR_ars_ is a common mechanism for control of *ars* operons and gene islands involved in resistance to arsenic-containing antibiotics in bacteria. Chemical warfare using arsenic containing compounds appears to be of ancient origin in microbes. Whether MarR_ars_-dependent is of ancient or relatively more recent origin remains to be determined. This finding enriches our knowledge about the regulation of genes that confer bacterial resistance to a wide variety of arsenic and antimony compounds.

## MATERIALS AND METHODS

### Strains, plasmids and primers

Strains, plasmids and primers used in this study are listed in Table 1. *Achromobacter* sp. As-55 and the *ΔmarR*_*ars*_ mutant were cultured at 28 °C aerobically in R2A medium (34). *E. coli* AW3110 (DE3) (*Δars::cam* F2IN(*rrn-rrnE*) bearing plasmids was grown aerobically in low phosphate medium (35) at 37 °C supplemented with the required antibiotics. CV17-Zero Background pTOPO-Blunt Simple Cloning Kit was purchased from Aidlab Biotechnologies Co., Ltd (Beijing, China) for construction of deletions. Plasmids pACYC184-P*marR*_*ars*_-*gfp* and pBAD-*AdmarR*_*ars*_ were constructed for biosensor assay (33). Primers of target genes used for RT-qPCR were designed using software Beacon designer 8.1.

**TABLE 1.**
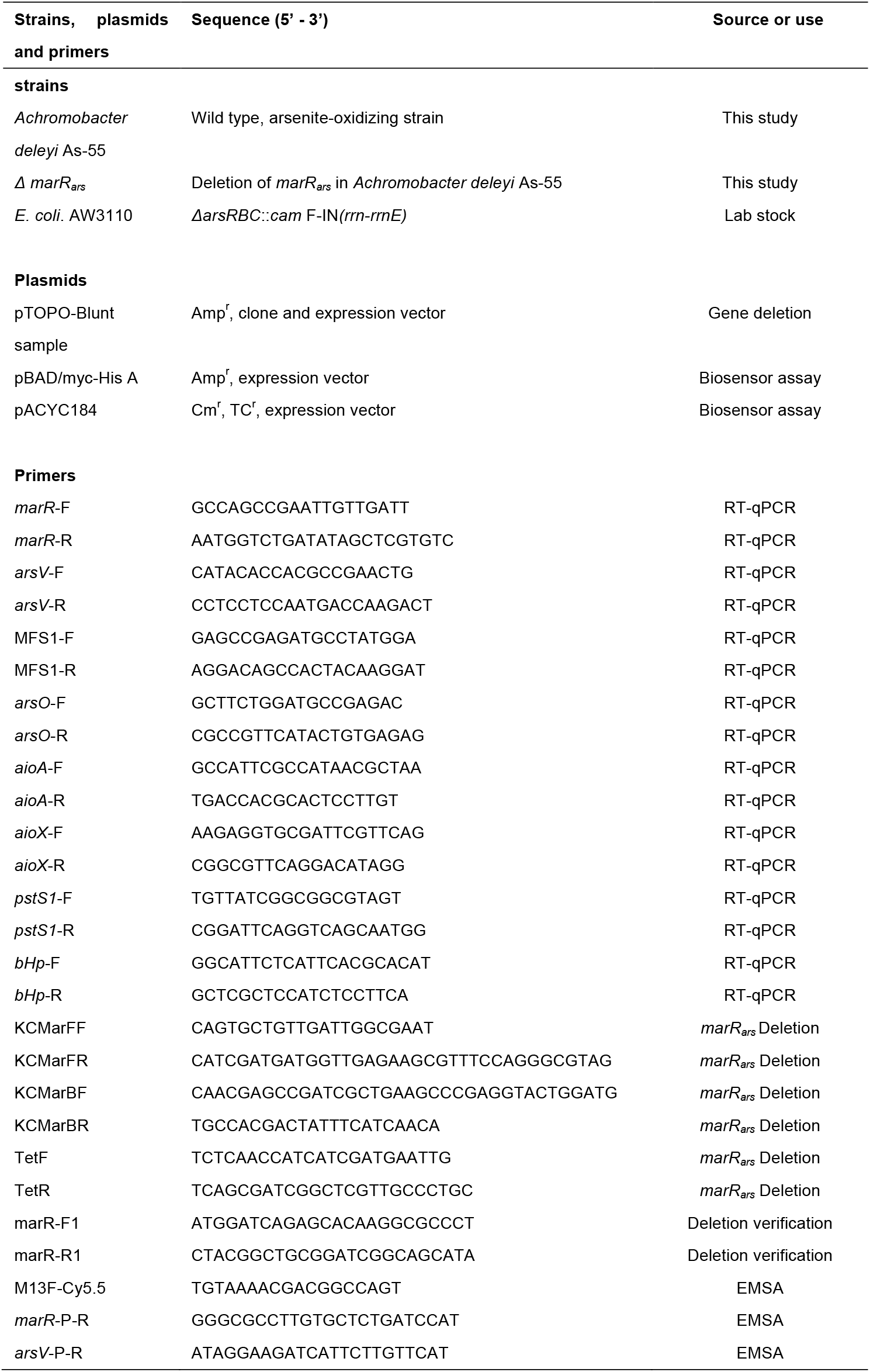
Strains, plasmids and primers used in this study.

### Distribution and sequence alignment of MarR_ars_ and phylogenetic analysis

The genome of *Achromobacter* sp. As-55 was sequenced using the Illumina Miseq platform. A putative *marR* gene was identified adjacent to an *ars*/*aio* operon in the draft genome of strain As-55 by functional gene annotation of Rapid Annotation using Subsystem Technology (RAST) (36). Acquisition of AdMarR_ars_ (MarR_*ars*_ protein of *Achromobacter* sp. As-55) homologous sequences was performed by searching a list of reference organisms or from the National Center for Biotechnology Information (NCBI) protein database using a BLASTP search (37). Multiple alignment of MarR homologs sequence was performed using Clustal Omega (http://www.ebi.ac.uk/Tools/msa/clustalo/). Phylogenetic analysis was performed to infer the evolutionary relationship among the representative *marR* adjacent or unrelated to *ars*/*aio* operons from various organisms. The phylogenetic tree was constructed using the Neighbor-Joining method with MEGA 6.0.1 (38). The statistical significance of the branch pattern was estimated by conducting a 1000 bootstrap.

### Deletion of *marR*_*ars*_ in *Achromobacter* sp. As-55

Homologous recombination was used to delete *marR*. In detail, two primer pairs of KCMarFF/KCMarFR and KCMarBF/KCMarBR were used to amplify the upstream and downstream sequence of *marR* respectively with genomic DNA of strain As-55 as template. Primers TetF and TetR were used to amplify the tetracycline resistant gene sequence (Table 1). Then the three sequences were assembled with primers KCMarFF and KCMarBR by overlap PCR. The assembled sequence contained the upstream sequence of *marR*, tetracycline resistant gene sequence and the downstream sequence of *marR*. The PCR product of the assembled sequence was purified with a DNA Fragment Purification Kit (Takara, Dalian of China) and eluted with ddH_2_O, then 10 μl of the purified sequence was added into 100 μl of competent cell of strain As-55, electroporation used a Gene Pulser Xcell™ (BIO-RAD) with 0.1 cm electroporation cuvettes (Cat: 1652083, BIO-RAD) at 25 μF, 200 Ohm and 1800 V. Agar plates of lysogeny broth medium (39) containing 15 mg L^-1^ of tetracycline were used for selecting positive transformants. Primers of *marR*-F1 and *marR*-R1 were used for *marR* deletion verification.

### Electrophoretic mobility shift assays

The DNA fragments of the *marR*_*ars*_ promoter and *arsV* promoter were amplified using two pairs of M13FCy5.5/*marR*-P-R and M13FCy5.5/*arsV*-P-R. All reaction mixtures were incubated at room temperature at dark condition in EMSA/Gel-Shift Binding Buffer (5X) (poly(dI-dC), DTT, glycerol, EDTA , NaCl , MgCl_2_ and Tris) for 20 min. Before being loaded onto a 6% PAGE gel, the binding solution was mixed with EMSA/Gel Shift Loading Buffer. After 2∼3 h of running at 60 V in 0.5 X TBE buffer, the gels were exposed in an imaging system (ODYSSEY CLx).

### Metalloid resistance assays

The *Achromobacter* sp. As-55 *arsV* gene was cloned and expressed in arsenic-hypersensitive strain *E. coli* AW3110 (*ΔarsRBC*). For metalloid resistance assays in liquid media, AW3110 cells bearing vector plasmid pTOPO or pTOPO-*arsV* were grown overnight with shaking at 37 °C in LB containing 100 mg L^-1^ Amp. The overnight cultures were inoculated into fresh LB medium with 100 mg L^-1^ Amp containing various concentrations of metal(loid)s and incubated at 37 °C with shaking for 24 h. The growth conditions were estimated using absorbance at OD_600_ nm. For metalloid resistance assays with wild type As-55 and *marR*_*ars*_ mutant *ΔmarR*_*ars*_, cells were streaked on R2A solid media containing different concentrations of metal(loid)s (As(III), Sb(III), Rox(III), Rox(V), Pb(II), Cu(II), Zn(II)).

### Total RNA extraction and RT-qPCR

A single colony of both strain As-55 and of mutant Δ*marR*_*ars*_ was incubated in R2A medium overnight. The cultures were diluted to an A_600nm_ of 0.01 into 30 mL of fresh R2A medium. 0.2 and 2 mM As(III) or 0.05 and 0.2 mM Sb(III) were added when the A_600nm_ reached 0.5, with no metal addition used as control. After incubation for 2 h, 1.5 ml or cells were harvested by centrifugation at 12,000 rpm for 2 min. Total RNA were extracted using a Trizol method (34), according to the manufacturer’s instructions. The RNA concentrations were quantified using a BioDrop Spectrophotometer (Biochrom Ltd, UK) and were diluted to appropriate concentrations before reverse transcription. cDNA was prepared by reverse-transcription PCR using HiScript^®^ III RT SuperMix for qPCR (+gDNA wiper) (Vazyme #R323, China). Briefly, gDNA contaminants present in the RNA samples were removed by treatment with 4 × gDNA wiper for 2 min at 42 °C, followed by reaction of 5 × HiScript III qRT SuperMix^a^ by following a program of 37 °C for 15 min and 85 °C for 5 sec. Quantitative real-time PCR was performed by using the QuantStudio 6 Flex Real-Time PCR System (Thermo Fisher Scientific, USA) with cDNA as the template. 16S rRNA of As-55 was used as an endogenous control. The relative expression was quantified according to the method of 2^−ΔΔCT^ (40).

### Construction of an AdMarR_ars_ homology model

The homology model of AdMarR_ars_ was constructed using the fully automated protein structure homology modeling server SWISS-MODEL (41) (http://swissmodel.expasy.org/). Model quality was estimated based on the QMEAN scoring function. The model was built using the structure of MarR from *Methanosarcina mazei* Go1 (PDB ID: 3S2W) as a template, the remainder was built using MODELLER without template with lower confidence. The sequence similarity and identity between the model and template are 30.0 and 20.9 %, respectively. The SWISSMODEL built residues from 10 to 148. The remaining residues from 149 to 163 were built using MODELLER program in CHIMERA software. PyMOL v1.6 was used to visualize the structural models (42) (https://www.pymol.org/citing).

### Mutagenesis of cysteine residues

Mutations in AdMarR_ars_ were introduced by site-directed mutagenesis using QuikChange II Site-Directed Mutagenesis Kit (Agilent Technologies, Santa Clara, CA). The mutagenic oligonucleotides used for both strands and the respective changes introduced (underlined) are as follows: C36SF: 5’-CCGCGATCGTGATCGCATT<u>AGC</u>TGCTATGACG-3’ and C36SR: 5’-CGTCATAGCA<u>GCT</u>AATGCGATCACGATCGCGG-3’; C37SF: 5’-ATCGTGATCGCATTTGC<u>AGC</u>TATGACGTTTCGGTA-3’ and C37SR: 5’-TACCGAAACGTCATA<u>GCT</u>GCAAATGCGATCACGAT-3’; C157SF: 5’-TGTCGCCTCCACA<u>AGT</u>GCTGCCGATCC-3’ and C157SR: 5’-GGATCGGCAGC<u>ACT</u>TGTGGAGGCGACA-3’. Each mutation was confirmed by commercial DNA sequencing (Sequetech, Mountain View, CA).

### Plasmid construction and assay of AdMarR_ars_ substrates binding *in vivo*

AdMarR_ars_ transcriptional activity was estimated from inducer-responsive biosensor activity measured by *gfp* expression (36). A *marR* gene corresponding to the mRNA sequence of the gene for AdMarR_ars_ (QVQ28260.1) in NCBI (CP074375.1) was chemically synthesized with 5’ *Nco*I and 3’ *SalI* sites and with codon optimization for expression in *E. coli* (GenScript, NJ, USA) and subcloned into expression vector pBAD/myc-His A (Invitrogen, Carlsbad, CA, USA) that produces a fusion six-histidine tag at the end. The *AdmarR*_*ars*_ promoter was chemically synthesized and subcloned into expression vector pACYC184 (NEB, United States), generating plasmid pACYC184-P*marR*_*ars*_-*gfp* (Fig. S6A). All the constructs were confirmed by DNA sequencing (Sequetech, Mountain View, CA). Cultures of the biosensor (*E. coli* strain AW3110 bearing plasmids pBAD-*AdmarR*_*ars*_, where the *marR*_*ars*_ gene is under control of the arabinose promoter, and pACYC184-P*marR*-*gfp*, where the *marR*_*ars*_ promoter is fused to a *gfp* gene) were grown to mid-exponential phase in low phosphate medium at 37 °C with 100 µg mL^-1^ ampicillin and 34 µg mL^-1^ chloramphenicol with shaking. Glycerol (0.5%) was added for constitutive expression of *gfp*. The *AdmarR*_*ars*_ gene was induced by addition of 0.2% arabinose for 5 h. Derepression was generated by simultaneous addition of arabinose and arsenicals for 5 h. Cell densities were normalized by dilution or suspension to the same A_600nm_, and expression of *gfp* was assayed from the fluorescence of cells using a Photon Technology International spectrofluorometer with an excitation wavelength. The GFP induction condition was shown in Fig. S6B.

## SUPPLEMENTAL MATERIALS

**SUPPLEMENTAL FILE 1**, PDF file, 0.1 MB.

## ACKNOWLEDGMENTS

This work was supported by the Natural Science Foundation of China (31770123), the Natural Science Foundation of Fujian province (2018J01668) to C.R., NIH grants R35 GM136211 and R01 GM55425 to B.P.R and the Natural Science Foundation of China (41967023) to J.C. We also acknowledge Researchers Supporting Project (RSP-2021/205), King Saud University, Riyadh, Saudi Arabia.

